# Geometric inference of cross-species transcriptome correspondence using Gromov-Wasserstein optimal transport

**DOI:** 10.1101/2025.10.23.684069

**Authors:** Yuya Tokuta, Tomonori Nakamura, Kohei Fujiwara, Masanori Imamura, Masahiro Nagano, Mitinori Saitou, Yusuke Imoto, Yasuaki Hiraoka

**Affiliations:** Institute for the Advanced Study of Human Biology, Kyoto University, Yoshida-konoe-cho, Sakyo-ku, Kyoto 606-8501, Japan; Department of Anatomy and Cell Biology, Graduate School of Medicine, Kyoto University, Yoshida-konoe-cho, Sakyo-ku, Kyoto 606-8501, Japan; Hakubi Center, Kyoto University, Yoshida-honmachi, Sakyo-ku, Kyoto 606-8501, Japan; Molecular Biology Section, Center for the Evolutionary Origins of Human Behavior, Kyoto University, Inuyama, Aichi 484-8506, Japan; Department of Medical Neuroscience, Graduate School of Medical Sciences, Kanazawa University, 13-1 Takara-machi, Kanazawa, Ishikawa 920-8640, Japan; Sapiens Life Sciences, Evolution and Medicine Research Center, Kanazawa University, 13-1 Takara-machi, Kanazawa, Ishikawa 920-8640, Japan; Department of Biological Engineering, Massachusetts Institute of Technology, 77 Massachusetts Avenue, Cambridge, MA 02139, USA; Center for iPS Cell Research and Applications, Kyoto University, 53 Kawahara-cho, Shogoin, Sakyo-ku, Kyoto 606-8507, Japan; Inamori Research Institute for Science, Inamori Foundation, 620 Suiginya-cho, Shimogyo-ku, Kyoto 600-8411, Japan; Center for Advanced Intelligence Project, RIKEN, 1-4-1 Nihonbashi, Chuo-ku, Tokyo 103-0027, Japan

**Keywords:** cross-species analysis, Gromov-Wasserstein optimal transport, gene correspondences, transcriptome data, bulk RNA-seq, scRNA-seq

## Abstract

Sequence homology reveals evolutionary gene correspondences, but identifying functionally corresponding genes within specific cell types or during cell fate specification remains non-trivial. Here, frameworks that translate insights from model organisms into other species would accelerate wet-lab studies, so we develop Species-OT, a cross-species transcriptome analysis framework using Gromov-Wasserstein optimal transport, which quantitatively compares the geometry of transcriptome distributions. Given a pair of bulk or single-cell RNA-sequencing datasets, Species-OT returns a gene-to-gene correspondence capturing probabilistic alignments of regulatory roles, and a transcriptomic distance quantifying overall divergence. Applied pairwise, Species-OT yields a transcriptomic discrepancy array and a hierarchical clustering tree analogous to a phylogenetic tree. We validate Species-OT using bulk RNA-seq data for cell fate specification from pluripotent stem cells in mice, monkeys, and humans and scRNA-seq data from pluripotent stem cells of six mammalian species. Species-OT identifies gene correspondences within cell fate specification, while transcriptomic discrepancies recapitulate expected species relationships.

## INTRODUCTION

Mammalian genomes and their evolutionary changes have been extensively studied. For example, the DNA sequences of humans and chimpanzees differ by about 1–2%, while approximately 85% of the protein-coding genes in the human and mouse genomes are conserved at the sequence level [1, 2, 3]. The concept of sequence homology has been formalized to describe the evolutionary relationships between genes derived from common ancestors, classifying those separated by speciation, which often retain similar biological functions, as orthologs, and those separated by gene duplication events as paralogs [4]. Paralogs can be further divided into inparalogs, which are paralogous genes that originated by duplication after a speciation event and therefore exist within a single species, and outparalogs, which are cross-species counterparts resulting from a duplication that occurred in a common ancestor before the speciation event [5, 6]. In contrast, regulatory elements such as promoters and enhancers exhibit pronounced species-specific divergence [7]. This has been largely due to the integration and selective co-option of distinctive transposable elements during species evolution, which results in the creation of species-specific gene regulatory networks (GRNs) [8]. Consequently, species-specific paralogs or evolutionarily distinct genes can play key roles in various cell fate specification processes and the horizontal transfer of findings from a model organism to other species does not necessarily guarantee an immediate success. For example, GRNs for cell fate specification in mammalian early development are divergent and consequently, the culture methods for early embryonic cells originally developed in mice have not been translatable to humans [9, 10].

Active wet-lab experiments have elucidated the functions of individual genes in each species and identified key regulators of various cell fate specification processes [11]. Furthermore, recent AI-driven methods enable the identification of corresponding cell types across species through transcriptome-based alignment using sequence homology and its extensions that incorporate protein sequence similarity and gene expression profiles [12, 13]. However, identifying functionally corresponding genes within specific cell types or during cell fate specification from the perspective of their regulatory roles, remains a nontrivial challenge. Thus, developing an approach that bridges the gap between omics datasets collected from model organisms and other species would greatly accelerate the transfer of biological knowledge across species.

To design such an approach, we pose our central question as “how can we identify gene correspondences in specific cell types or during cell fate specification processes from the perspective of regulatory roles across species?” We can hypothesize that the core parts of underlying GRNs of conserved biological processes in different species have similar architectures, while individual gene factors of the GRNs may be swapped. Based on this hypothesis, we propose a strategy to tackle that biological question by comparing the structure of GRNs and their gene factors among multiple species, with a particular emphasis on the subnetworks consisting of transcription factor genes as their core parts. Given our limited knowledge of gene correspondences from the perspective of regulatory roles across species, it is necessary to consider correspondences between GRNs that involve many transcription factors and regulatory elements while minimizing potential selection bias. However, direct construction and comparison of large-scale GRNs across species are impractical, as the associated computational costs result in a combinatorial explosion [14]. Moreover, if we attempt to account for all the diverse factors, their complexity may obscure the simpler common features. Therefore, we conceptually simplify the target and seek to uncover the insights that transcriptome data can reveal, focusing on the observation that the geometry of transcriptome distributions inherits similar geometric structures from the underlying GRNs and tackle that biological question by developing a method to compare the geometry of transcriptome distributions obtained from different species.

One mathematical approach for geometrically comparing distributions is to use optimal transport, which quantifies the difference between two probability distributions by minimizing the cost of transporting one into the other. Optimal transport has recently gained traction in omics data analysis: for example, Waddington-OT [15] applies unbalanced optimal transport to single-cell RNA-sequencing (scRNA-seq) time-course data to infer differentiation trajectories at the single-cell level. A related formulation, Gromov-Wasserstein optimal transport, extends this concept to distributions in different metric spaces by defining a transport cost based on the discrepancy between their geometric representations, such as distance matrices. For instance, applying Gromov-Wasserstein optimal transport to pairwise combinations of transcriptome, tissue, and epigenome data has enabled the integration of multi-omics datasets [16, 17, 18].

We present Species-OT, a cross-species transcriptome analysis method that applies Gromov-Wasserstein optimal transport to transcriptome datasets from different species. Species-OT is applicable to both bulk RNA-seq and scRNA-seq data without requiring pre-alignment of gene lists or cell types among species, leveraging the space-free property of Gromov-Wasserstein optimal transport. It computes a gene-to-gene correspondence that captures probabilistic alignments of regulatory roles of genes across species and a transcriptomic distance that quantifies overall divergence. These are biological reinterpretations of the entropic optimal transport plan and the entropic Gromov-Wasserstein distance, which are the mathematical outputs of Gromov-Wasserstein optimal transport, in light of the biological hypothesis described above. Unlike traditional sequence homology-based approaches, Species-OT focuses on transcriptome profiles rather than DNA sequences, enabling the identification of gene correspondences from the perspective of their regulatory roles. Moreover, when extended to multiple species, Species-OT generates a transcriptomic discrepancy array that captures differences in transcriptome distributions across species, and its hierarchical clustering tree provides a visualization analogous to a phylogenetic tree, redefining inter-species relationships at the transcriptomic level.

To demonstrate the power of Species-OT, we apply it to bulk RNA-seq data for the specification of the germ cell fate from pluripotent stem cells in mice, monkeys, and humans, as well as to scRNA-seq data from pluripotent stem cell lines of six mammalian species. We validate Species-OT by assessing its ability to recover biologically studied gene correspondences in human and mouse germ-cell lineage specification, out of all the computed gene correspondences. Furthermore, we present an example of a transcriptomic discrepancy tree among mammalian species that deviates from the phylogenetic species relationships, utilizing the discrepancy measuring inter-species differences influenced by culture conditions. Our approach overcomes previous limitations in identifying gene correspondences within specific cell types or during cell fate specification from the perspective of regulatory roles, achieving higher resolution than many-to-many paralog correspondences by leveraging geometric comparisons of transcriptomes, thereby revealing species-specific differences in conserved biological processes.

## RESULTS

### Overview of Species-OT

The biological hypothesis behind Species-OT (Figure 1A) is that gene expression in conserved cell types or during cell fate specification of distinct species should be governed by GRNs with conserved core architectures, while individual gene factors comprising the GRNs may be swapped. This is because gene functions may diverge at speciation, or distinct paralogs may replace each other, as in the case of human *EOMES* and mouse *T*, each of which acts upstream in the in vitro recapitulation of germ cell lineage specification, referred to as primordial germ cell-like cell (PGCLC) induction [11]. GRNs with similar core architectures cause analogous key gene expression patterns, resulting in similar *transcriptome distributions* formed by filtered transcriptomes. However, because the orders and labels of cells and genes differ across species, these distributions may be misaligned in orientation. In other words, genes with equivalent regulatory roles occupy the same relative positions within each distribution but differ in their absolute coordinates. To identify gene correspondences across species, we therefore align these shapes and compare gene positions on the aligned distributions.

**Figure 1.**
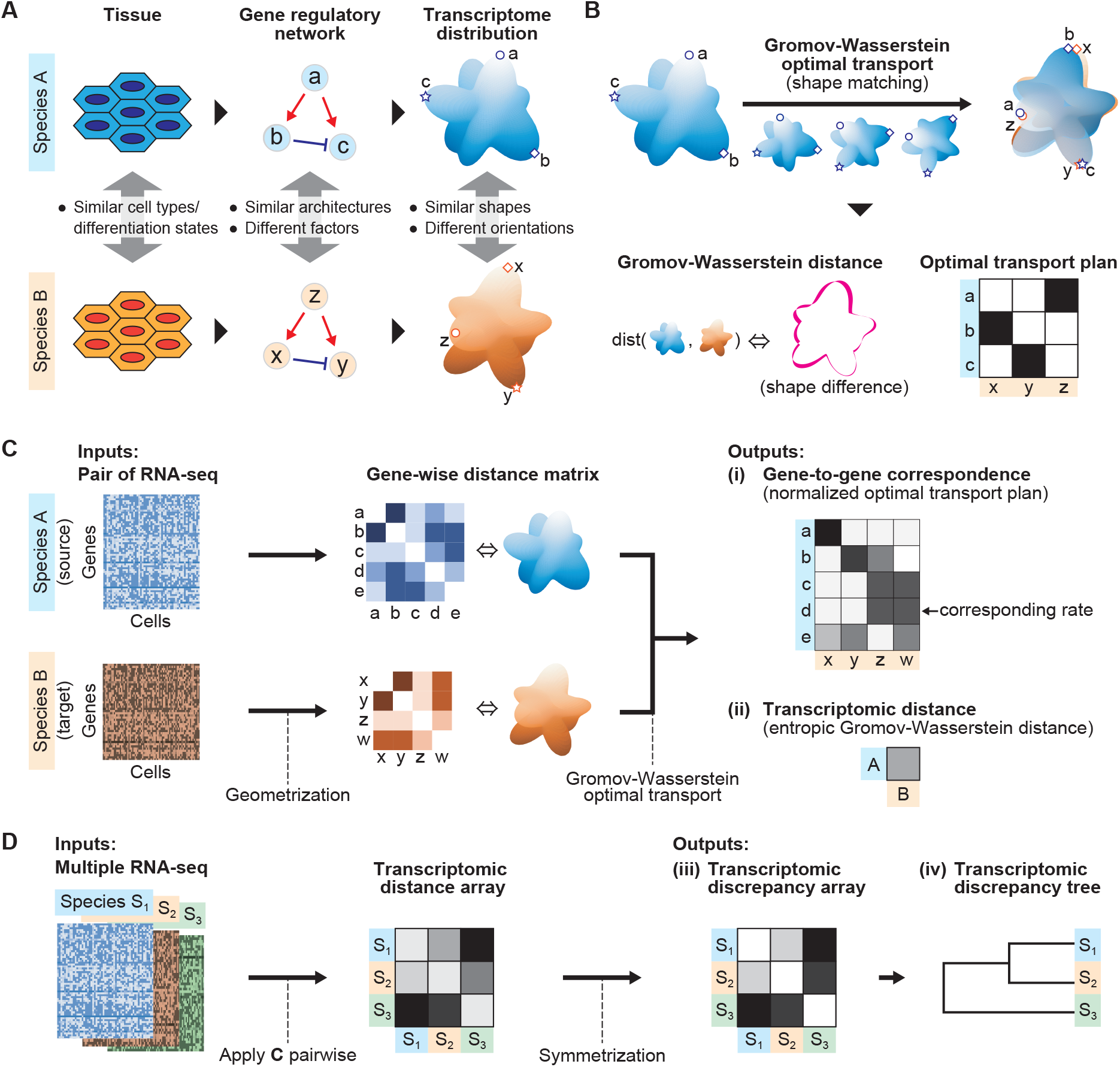
Overview of the Species-OT framework. **A.** Biological hypothesis. Conserved cell types or cell fate specification processes share similar core GRN architectures, with possible divergence in gene factors, results in similarly shaped transcriptome distributions with varying orientation and scale. **B**. Illustration of Gromov-Wasserstein optimal transport (GWOT). GWOT minimizes the sum of the products of the differences between lengths of corresponding edges and transported masses, visualized as the shape difference in transcriptome distributions. **C**. Species-OT workflow for a pair of RNA-seq datasets *X*^A^, *X*^B^. First, the datasets are geometrized into gene-wise distance matrices *M* ^A^, *M* ^B^. GWOT then yields (i) gene-to-gene correspondence 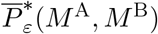, a normalized transport plan encoding correspondence probabilities of genes between species, and (ii) transcriptomic distance *D*_2,*ε*_(*M* ^A^, *M* ^B^), an entropic Gromov-Wasserstein distance quantifying the geometric divergence of their transcriptome distributions. **D**. Extension to multiple species S_1_, … S_*K*_. Pairwise application of Species-OT produces an array of transcriptomic distances 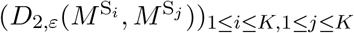. Symmetrization of this array yields (iii) transcriptomic discrepancy array 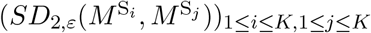, which can be visualized as a dendrogram, (iv) transcriptomic discrepancy tree.

To achieve this, we employ Gromov-Wasserstein optimal transport for geometric comparison of transcriptome distributions (Figure 1B). Gromov-Wasserstein optimal transport finds the optimal alignment between two distributions by quantifying differences in their geometric representations, such as distance matrices. Since a distance matrix is invariant under rotation and translation, this method captures only shape differences. It returns the Gromov-Wasserstein distance, which measures the total geometric gap between distributions, and the optimal transport plan, which encodes pairwise correspondences between points in the aligned distributions. Although the Gromov-Wasserstein distance is insensitive to orientation changes, the optimal transport plan holds information about evolutionary substitutions of GRN factors. Specifically, substituted genes are represented as permutations of the columns in the optimal transport plan. Consequently, the optimal transport plan derived from the two species’ transcriptome distributions directly reflects gene correspondences. The precise mathematical formulation is described in the method details section.

We present Species-OT, a cross-species transcriptome analysis method built on Gromov-Wasserstein optimal transport (Figure 1C). Species-OT takes as input a pair of transcriptome data matrices, either scRNA-seq or bulk RNA-seq. It first preprocesses the data and then converts each matrix into a gene-wise distance matrix in the *geometrization* step. Next, it solves the entropically regularized Gromov-Wasserstein optimal transport problem between these distance matrices to find the optimal transport plan that minimizes the cost of their geometric differences. Here, entropic regularization is a regularization of the cost used for stabilizing computation. Finally, Species-OT outputs (i) the *gene-to-gene correspondence*, a normalized transport plan encoding correspondence probabilities between genes of the two species, and (ii) the *transcriptomic distance*, which quantifies the geometric divergence of their transcriptome distributions. The entry of gene-to-gene correspondence, called the *corresponding rate*, shows a probability of correspondence from the gene of the source species in row to the gene of the target species in column.

Further, the framework of Species-OT can be extended to multiple species (Figure 1D). Taking multiple RNA-seq datasets, Species-OT handles each pair of input datasets and provides us with the *transcriptomic distance array*. Applying Sinkhorn-inspired symmetrization to the transcriptomic distance array to correct for its ordering bias results in (iii) the *transcriptomic discrepancy array*. Finally, hierarchical clustering of this array produces (iv) the *transcriptomic discrepancy tree*, a dendrogram visualization of inter-species transcriptomic relationships, akin to a phylogenetic tree.

### Gene-to-gene correspondences between mice and humans for PGCLC induction

This section demonstrates gene correspondences between mouse and human PGCLC induction processes as an example of analyzing cell fate specification processes of multiple species represented by bulk RNA-seq datasets. We validate the computed correspondences by comparing them with biologically studied references. Saitou et al. [19] reviewed previously characterized evolutionarily distinctive transcriptional and signaling programs during PGCLC induction based on experimental and evolutionary analyses (Figure 2) [11, 20]. We assess whether Species-OT can accurately recover these biologically identified correspondences.

**Figure 2.**
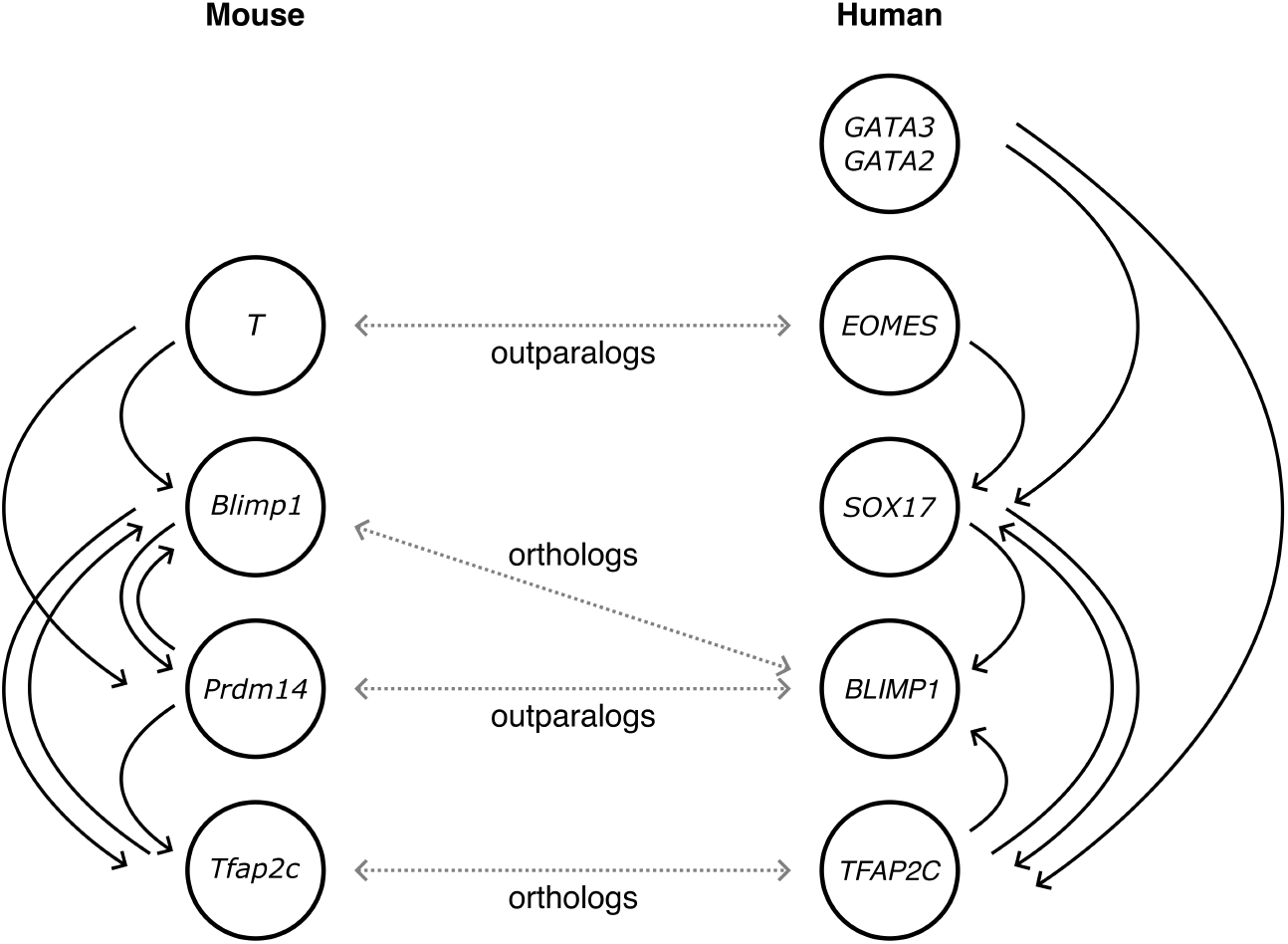
Biological correspondences between key genes involved in PGCLC induction in mice and humans [11, 19, 20]. Adapted from the third figure, *Signaling and transcriptional network for PGCLC induction in mice and humans*, in the review paper by Saitou and Hayashi [19]. Arrows in black indicate regulatory relationships among signaling and transcription factors. Genes connected by a dotted arrow in gray correspond to each other from the perspective of their regulatory relationships within the signaling and transcriptional networks and homological labels are added [21].

For the key genes during PGCLC induction (Figure 2), *T, Blimp1, Prdm14*, and *Tfap2c* for mouse and *GATA2, GATA3, EOMES, SOX17, BLIMP1*, and *TFAP2C* for human, we investigated the top 10 corresponding genes for each based on the computed corresponding rates, the entries of the gene-to-gene correspondences (Figure 3).

**Figure 3.**
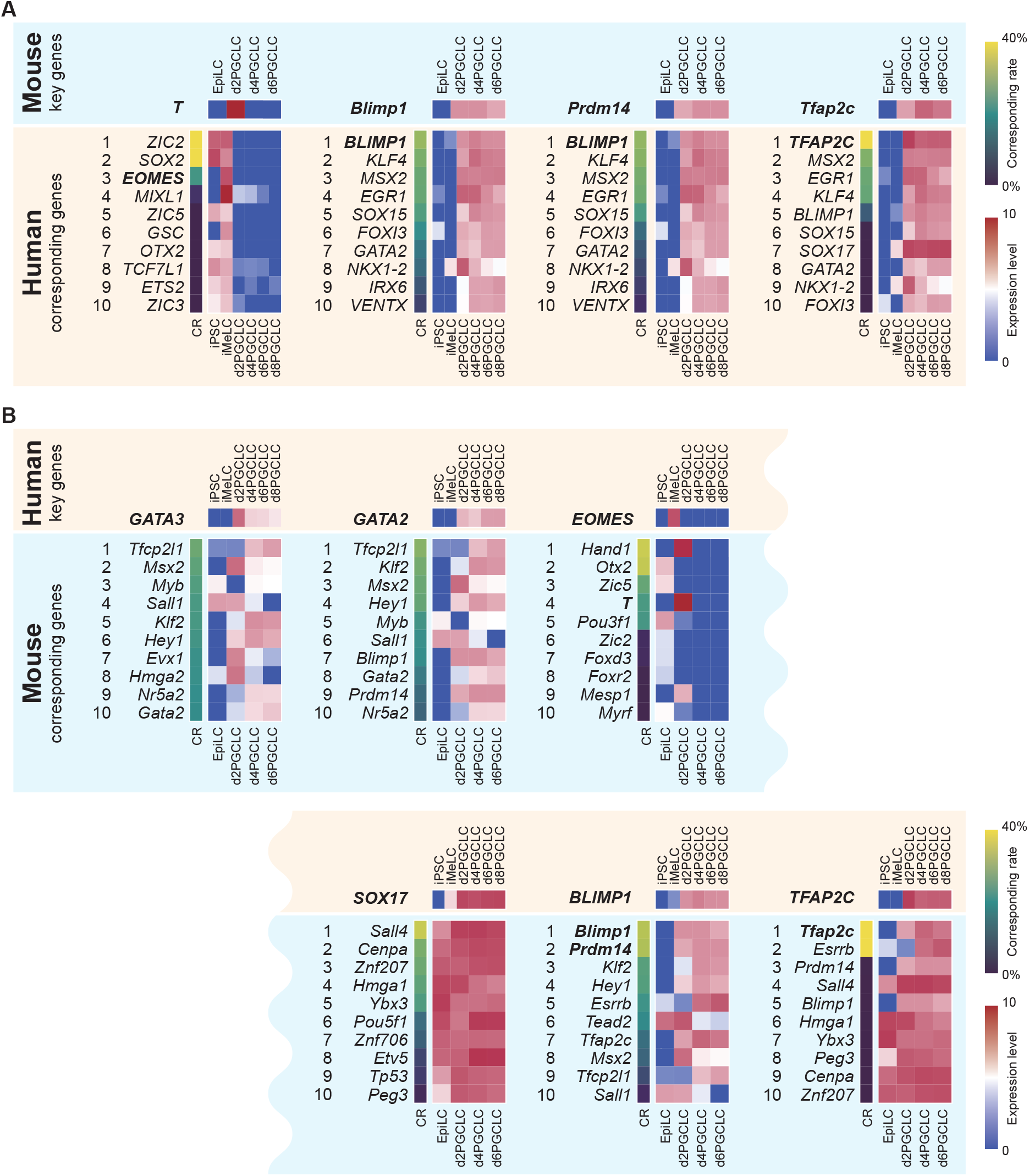
Comparison of key genes and their top-ranked corresponding genes between mouse and human PGCLC induction.**A.** Corresponding rates (CR) and expression profiles of four mouse key genes alongside the top 10 corresponding human genes. **B**. Corresponding rates (CR) and expression profiles of six human key genes alongside the top 10 corresponding mouse genes. The key genes and their biologically corresponding genes, shown in Figure 2, are written in bold style. The expression profiles are averaged across two replicates at each time-point.

For the mouse key genes, Species-OT successfully detected all biologically corresponding human genes among approximately 1,000 genes that remained after preprocessing (Figure 3A). In particular, for mouse *Blimp1, Prdm14*, and *Tfap2c*, their biologically corresponding human genes, *BLIMP1, BLIMP1*, and *TFAP2C*, respectively, were ranked first, whereas for mouse *T*, one of its paralogs human *EOMES* with the known important role in germ cell specification was ranked third. Likewise, for human *BLIMP1*, its biologically corresponding mouse genes, both *Blimp1* and *Prdm14* were ranked first and second, for human *TFAP2C*, its biologically corresponding mouse gene *Tfap2c* was ranked first, and for human *EOMES*, its biologically corresponding mouse paralog *T* was ranked fourth (Figure 3B). Thus, Species-OT successfully identified the known corresponding mouse key genes to the human key genes, highlighting the robustness of swapping source and target species datasets. We can further refine these computational results of gene correspondences, which we will address in the discussion section.

Next, we turn our attention to previously uncharacterized corresponding genes. Human *SOX17* is a specifier of human PGCLC fate [22], and its top corresponding gene, mouse *Sall4*, facilitates mouse PGCLC specification by repressing somatic program genes [23]. Thus, both genes are essential for PGCLC specification in their respective species, and their relationship may represent an evolutionarily distinct correspondence. Furthermore, despite some differences in expression dynamics, mouse *Tfcp2l1* was ranked first among correspondences to both human *GATA3* and *GATA2*, which, together with *SOX17* and *TFAP2C*, drive the human PGCLC program [20]. This observation may reflect their analogous positions within the signaling and transcriptional network (Figure 2). Therefore, Species-OT has the potential to uncover hidden biological insights by closely examining computed gene-to-gene correspondences.

### Cross-species analysis of PGCLC induction processes using bulk RNA-seq data

We validated Species-OT across multiple species by including bulk RNA-seq datasets of the PGCLC induction system from macaque, in addition to those from human and mouse. The resulting transcriptomic discrepancy array accurately reflects biological expectation: human and macaque cluster closely, while mouse is more distant (Figure 4A). The transcriptomic discrepancy tree, a dendrogram of these array values, further clarifies their relationships, resembling a phylogenetic tree (Figure 4B). Indeed, the simplified phylogeny corroborates our inferred relationships (Figure 4C). However, the ratio of transcriptomic discrepancies between the human–macaque branch and mouse (0.012 : 0.043 *≈* 1 : 3.58) differs from the corresponding divergence times in the phylogeny (30 : 80 *≈* 1 : 2.67). This difference likely reflects fundamental differences between transcriptomic discrepancy, capturing cellular-level expression divergence, and evolutionary distance derived from the molecular analysis of the DNA sequences.

**Figure 4.**
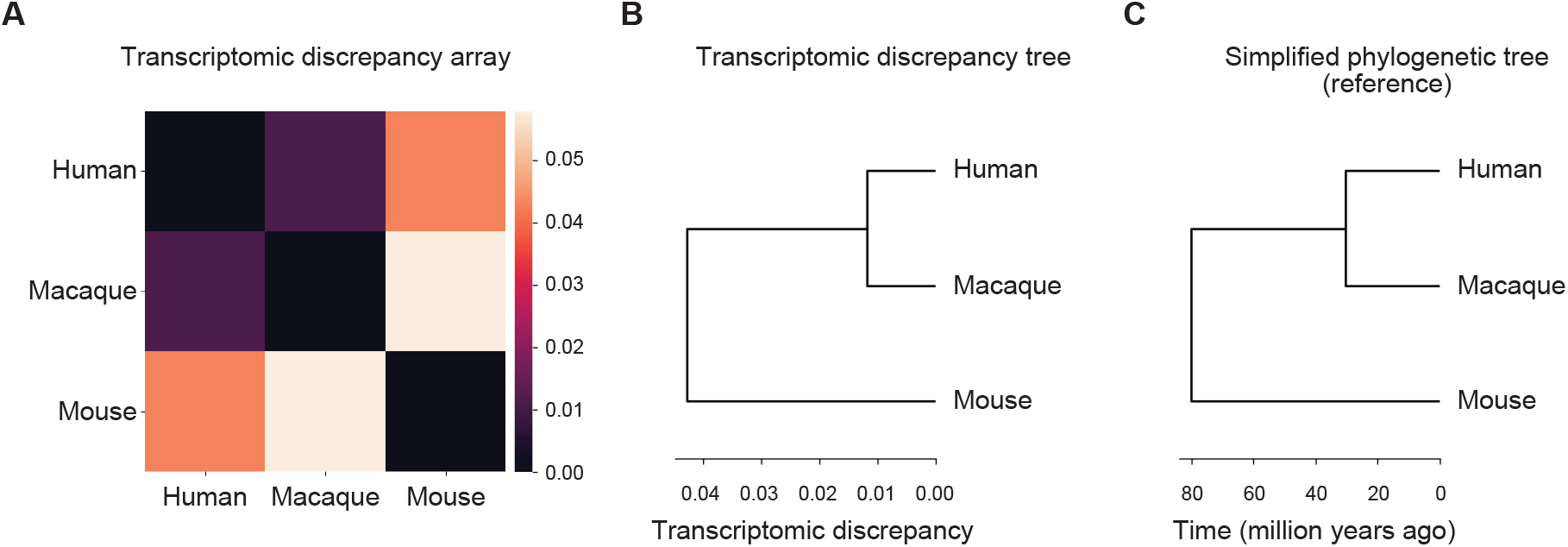
Validation results of transcriptomic discrepancy array and tree.**A.** The transcriptomic discrepancy array of bulk RNA-seq data from the three species. Here, the rows represent the source species and the columns represent the target species, from which the bulk RNA-seq data were collected. **B**. Transcriptomic discrepancy tree of three species derived from the transcriptomic discrepancy array using hierarchical clustering. **C**. Simplified phylogenetic tree for the corresponding three species, made by referring to sequence homological studies [24, 25].

### Cross-species analysis of pluripotent stem cells using scRNA-seq data

We applied Species-OT to scRNA-seq data from six pluripotent stem cell lines: human, chimpanzee, gorilla, and orangutan iPSCs; macaque ESCs; and mouse epiblast-like cells (EpiLCs). Unlike the PGCLC induction data, these scRNA-seq profiles derive from a single cell type at one time-point, so Species-OT captures intrinsic cell type transcriptomic differences. The resulting transcriptomic discrepancy array reveals a pronounced separation of mouse from the primates, as well as subtler distinctions among primates (Figure 5A). Furthermore, the absolute values of the transcriptomic discrepancy array are lower than those observed for the PGCLC induction system in the previous section (Figure 4A), reflecting the relatively static nature of pluripotent stem cell lines.

**Figure 5.**
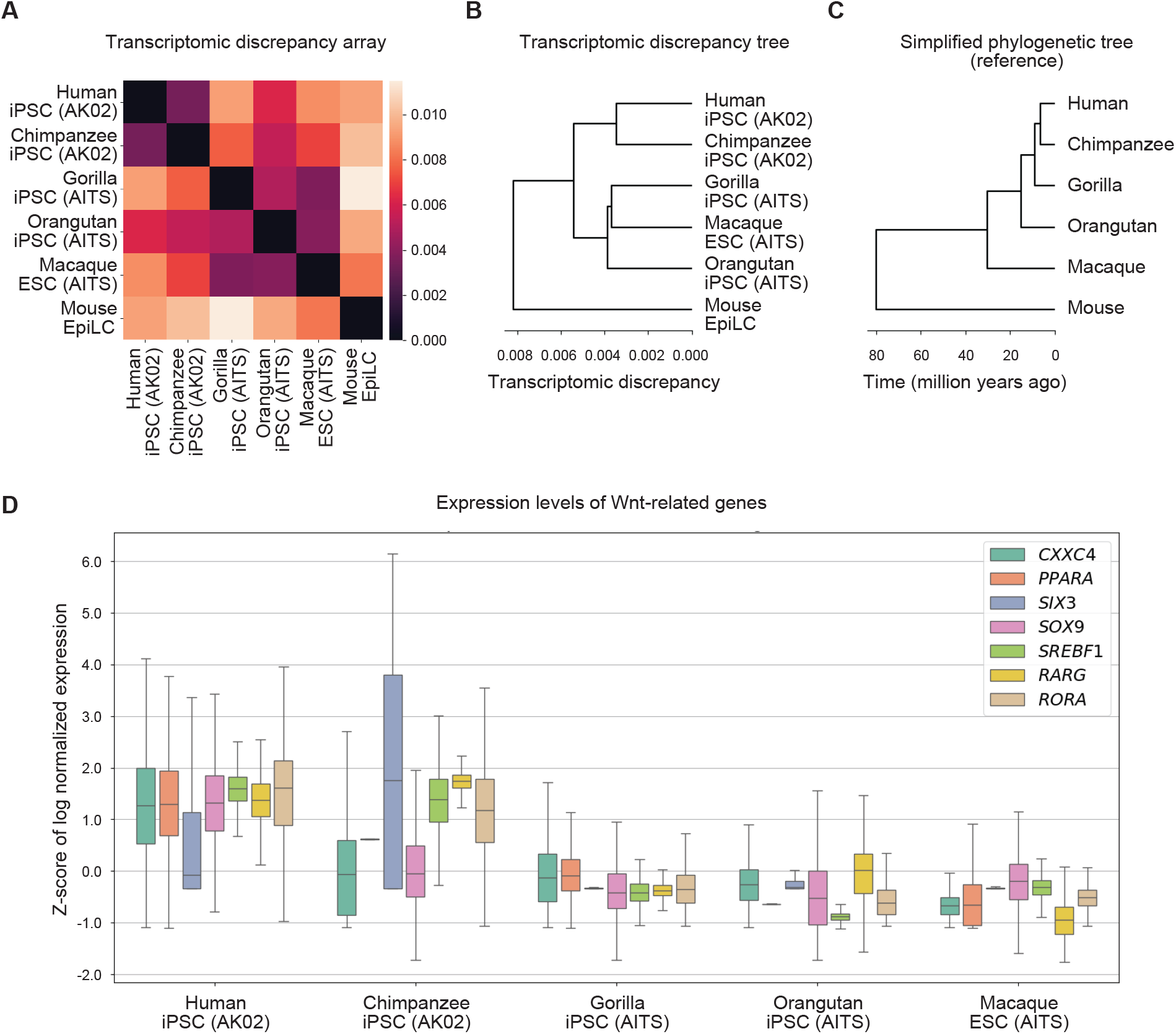
Validation results of transcriptomic discrepancy array and tree.**A.** The transcriptomic discrepancy array of scRNA-seq data from the six species. Here, the rows represent the source species and the columns represent the target species, from which the scRNA-seq data were collected. **B**. Transcriptomic discrepancy tree of six species derived from the transcriptomic discrepancy array using hierarchical clustering. **C**. Simplified phylogenetic tree for the corresponding six species, created by referring to sequence homological studies [24, 25]. **D**. Comparison of expression levels of downstream transcription factors to the Wnt signaling pathway across culture conditions. Boxplots show the Z-score of normalized log expression values for representative Wnt target transcription factor genes under two different culture conditions, AK02 and AITS. Each colored box represents one gene. The expression of these genes is generally higher in AK02, consistent with the known activation of Wnt signaling in this condition. In contrast, the AITS condition is associated with reduced expression of these genes, consistent with suppression of Wnt signaling pathway activity.

The corresponding transcriptomic discrepancy tree (Figure 5B) groups the primates into two clusters: human-chimpanzee and orangutan-gorilla-macaque, contrasting with the phylogenetic hierarchy (Figure 5C) and suggesting that cell state or culture condition differences may drive these patterns. Indeed, the clustering aligns with two culture media, AK02 versus AITS. These culture media, for example, differ in Wnt inhibition: AITS includes Wnt pathway inhibitors, which are associated with reduced expression of Wnt-responsive or Wnt-associated genes such as *CXXC4* and *PPARA*, whereas AK02 does not contain such inhibitors. We observe corresponding expression changes in these factors (Figure 5D). Additional variations in culture components, such as FGF signaling strength and feeder-cell dependence, also affect non-pluripotency gene expression, indicating that media differences, rather than species divergence, underlie the observed transcriptomic patterns. Thus, Species-OT accurately quantifies cell type transcriptomic differences without being confounded by species origin.

## DISCUSSION

We have developed Species-OT, a cross-species transcriptome analysis method using Gromov-Wasserstein optimal transport, which is a mathematical theory that quantifies the difference between probability distributions. For a pair of transcriptome datasets from distinct species, Species-OT computes a gene-to-gene correspondence capturing the probabilistic alignments of genes from the perspective of regulatory roles across species. For multiple transcriptome datasets, Species-OT generates a transcriptomic discrepancy array and a tree quantifying transcriptomic relationships among species. Applying Species-OT to bulk RNA-seq data collected at multiple time-points during mouse and human PGCLC induction processes, we identified biologically known ortholog correspondences (mouse *Blimp1* and human *BLIMP1*; mouse *Tfap2c* and human *TFAP2C*) as well as outparalog correspondences (mouse *T* and human *EOMES*; mouse *Prdm14* and human *BLIMP1*); and computed corresponding genes for all highly expressed transcription factor genes, including unexplored candidate pairs like mouse *Sall4* and human *SOX17*. Furthermore, Species-OT quantified differences in bulk RNA-seq time-course data during PGCLC induction in human, macaque, and mouse and in scRNA-seq data from pluripotent stem cells of six mammalian species, finding that human and macaque induction patterns are more similar to each other than to those of mouse, and that pluripotent stem cell lines cluster according to the evolutionary proximity influenced by culture conditions, formulating inter-species differences at the cell type level.

The computed gene-to-gene correspondences could be further refined by incorporating ad hoc biological observations. For example, all the key genes of both mouse and human are silent at the initial time-point (Figure 3), but the top two corresponding human genes *ZIC2* and *SOX2* to mouse gene *T* are significantly highly expressed in iPSCs before the onset of induction, making it questionable that these genes *ZIC2* and *SOX2* are biologically corresponding to the induction-critical mouse gene *T*. We could then exclude human *ZIC2* and *SOX2* as candidate genes corresponding to mouse *T*, and human *EOMES* could move up as the most corresponding human gene to mouse *T*. Similarly, we could exclude mouse *Otx2* and *Zic5* as corresponding genes for human *EOMES*, even though human *EOMES* and mouse *Otx2* pair might be biologically relevant despite their difference in early differentiation dynamics, with *Otx2* being low in naive pluripotent cells and upregulated upon priming, showing high expression on the opposite side of the PGCLC trajectory and in endodermal lineages [26]. Moreover, human *EOMES* and mouse *Hand1* pair could be regarded as an example of genes which are expressed in the corresponding timing with distinct roles, as their divergent lineage trajectories of mesoendodermal and amniotic progenitors during early development are known [27, 28].

Species-OT has the advantage of operating on geometric shapes extracted from transcriptome data, enabling the comparison of relative relationships among all genes of each species having distinct gene sets and varying cell types. By matching the shapes of transcriptome distributions formed by hundreds or thousands of genes, Species-OT yields comprehensive and quantitative metrics that differ fundamentally from traditional analyses of individual genes and their expression patterns. Consequently, Species-OT offers an alternative to homology-based or expression-pattern methods and broadens the range of compatible data.

A main limitation of Species-OT is that it cannot identify gene correspondences between cell types or processes without well conserved core parts of underlying GRNs. In our gene correspondence analysis, we successfully identified ortholog correspondences at the highest ranks without referring to their gene expression. On the other hand, for outparalog correspondences, we suggested excluding human *ZIC2* and *SOX2* as the corresponding genes for mouse *T*, and mouse *Otx2* and *Zic5* for the corresponding genes for human *EOMES*, based on their gene expression patterns. This implies that the core parts of underlying GRNs may differ slightly between mouse and human germ cell specification, owing to evolutionary changes associated with species divergence. Furthermore, there are differences between in vivo cell fate specification processes and their in vitro recapitulation models. Ideally, we aim to identify gene correspondences without relying on ad hoc biological insights. To further refine the computational quality of our approach in future work, we plan to account for variations in underlying GRNs arising from biological or experimental factors and to incorporate epigenomic data into our analysis.

In principle, the Gromov-Wasserstein optimal transport framework can be applied to any omics modality, including ATAC-seq [29], ChIP-seq [30], Hi-C [31], quantitative proteomics [32], and beyond, for both cross-species and other comparative analyses. However, such applications do not always yield meaningful biological insights, since each setting requires its own biological hypothesis to interpret the outputs. In this study, we hypothesized that equivalent cell types across species are governed by GRNs with similar core architectures, and that these GRNs give rise to comparably shaped transcriptome distributions. Under this hypothesis, matching shapes of transcriptome distributions using Gromov-Wasserstein optimal transport signifies aligning gene factors comprising GRNs, allowing us to biologically interpret both the resulting entropic optimal transport plan and the entropic Gromov-Wasserstein distance. Careful formulation of such hypotheses will be essential for extending Species-OT to other contexts.

In response to the central question of this study, “how can we identify gene correspondences in specific cell types or during cell fate specification processes from the perspective of regulatory roles across species?”, we offered an answer through both mathematical and biological approaches, utilizing transcriptome data collected from the target cell types or cell fate specification processes. Mathematically, we formulated a methodology to address the challenge of comparing datasets with differing gene sets and cell types by geometrically matching the shapes of transcriptome data using Gromov-Wasserstein optimal transport. Biologically, we interpreted the resulting entropic optimal transport plans as probabilistic alignments of genes from the perspective of regulatory roles across species, based on the hypothesis that transcriptome distributions inherit similar core architectures of conserved gene regulatory networks. Thus, by integrating mathematical rigor with biological insight, we resolved this question and demonstrated how such cross-disciplinary research can drive progress in tackling biological challenges, such as development of cross-species analysis. Species-OT, born from cross-disciplinary research, enables us to cross-map biological insights across species and will contribute to accelerating biological research.

## RESOURCE AVAILABILITY

### Lead contact

Requests for further information should be directed to and will be fulfilled by the lead contact, Yuya Tokuta.

### Materials availability

Great ape cells and molecular materials are available subject to the following regulations; institutional regulations and MTA in the Center for the Evolutionary Origins of Human Behavior (EHUB), Kyoto University, the Great Ape Information Network (GAIN), and international regulations according to the Convention on International Trade in Endangered Species of Wild Fauna and Flora (CITES).

### Data and code availability

Mouse PGCLC data were obtained from the NCBI Gene Expression Omnibus (GEO; accession GSE67259), as described in the experimental model and subject details section. Bulk RNA-seq data from the PGCLC induction systems of human and cynomolgus macaque, as well as scRNA-seq data from pluripotent stem cells of six mammalian species used in this study, will be available under GEO accession GSE307647 after publication of this manuscript. The processed transcriptome datasets will also be uploaded as supplemental files.

Species-OT is distributed as a Python package and can be installed via pip. The source code is open-source and available at https://github.com/yuyatokuta/Species-OT.

## ACKNOWLEDGMENTS

Yudai Yasui assisted in refactoring and publishing the source code online. Hanbo Wang provided valuable feedback on human and mouse PGCLC induction. We thank our colleagues for their helpful input on this study. We are grateful to Dr. Keisuke Okita for providing the orangutan iPSC line TobaK, the Single Cell Genome Information Analysis Core of WPI-ASHBi for their assistance with high-throughput DNA sequencing analysis, the Cooperative Research Programs of the Primate Research Institute (PRI), the Center for the Evolutionary Origins of Human Behavior (EHUB), Kyoto University, and the Great Ape Information Network (GAIN). This work was sponsored by JST CREST (JPMJCR2334 to Y.H. and JPMJCR24Q1 to Y.I.), JST Moonshot R&D (JPMJMS2021 to Y.H.), JST MIRAI (JPMJMI22G1 to Y.H.), JST PRESTO (JPMJPR2021 to Y.I.), JST FOREST (JPMJFR222X to Y.I.), JSPS Grant-in-Aid for Transformative Research Areas (A) (22H05107 to Y.H.), JSPS Grant-in-Aid for Scientific Research (C) (19K06864 to M.I.), the Cooperative Research Programs of the Primate Research Institute Kyoto University (2019-C-6, 2020-B-50, 2021-A-24 to M.S.).

## AUTHOR CONTRIBUTIONS

Y.T., T.N., M.S., Y.I., and Y.H. designed the study; Y.I. conducted the pilot study; Y.T. conducted the research; Y.T. developed the computational method; Y.T. analyzed the data; M.I. and K.F. established the great ape cell lines; T.N., K.F., and M.N. collected and set up the bulk or single-cell RNA-seq data; Y.T., T.N., Y.I., and Y.H. wrote the manuscript; and everyone reviewed the manuscript.

## DECLARATION OF INTERESTS

The authors declare no competing interests.

### Materials and methods

#### Method details

##### Input data

The bulk RNA-seq data were generated from human, macaque, and mouse cells in vitro culture. Human cells include iPSCs, incipient mesoderm-like cells (iMeLCs), and PGCLCs from day 2 to day 8 at four time-points (d2PGCLC–d8PGCLC). Macaque cells include ESCs and PGCLCs at four time-points. Mouse cells include epiblast-like cells (EpiLCs), which correspond more closely to the primed state of human iPSCs than mouse ESCs [33], and PGCLCs at three time-points. We prepared two replicates for each time-point. These are summarized in Table 1. Mouse PGCLC data were obtained from NCBI-GEO (GSE67259) as described in the experimental model and subject details section.

**Table 1:** Summary of the bulk RNA-seq data from human, macaque, and mouse.

|  | human | macaque | mouse |
| --- | --- | --- | --- |
| # genes | 22,180 | 20,583 | 14,800 |
| # bulk samples | 12 | 10 | 8 |
| cells (time-points) | iPSC | ESC | EpiLC |
|  | iMeLC | d2PGCLC | d2PGCLC |
|  | d2PGCLC | d4PGCLC | d4PGCLC |
|  | d4PGCLC | d6PGCLC | d6PGCLC |
|  | d6PGCLC | d8PGCLC | - |
|  | d8PGCLC | - | - |

The scRNA-seq data were generated from human, chimpanzee, gorilla, orangutan, macaque, and mouse pluripotent stem cells. We prepared human and chimpanzee iPSCs using the AK02 culture method [34], gorilla and orangutan iPSCs and macaque ESCs the AITS culture method, and mouse EpiLCs the standard method. These pluripotent stem cells serve as the initial phase of PGCLC induction, and they were collected at a single time-point. These are summarized in Table 2.

**Table 2:** Summary of the scRNA-seq data from human, chimpanzee, gorilla, orangutan, macaque, and mouse.

|  | human | chimpanzee | gorilla | orangutan | macaque | mouse |
| --- | --- | --- | --- | --- | --- | --- |
| # genes | 32,147 | 28,521 | 29,854 | 28,684 | 25,898 | 21,109 |
| # cells | 2,895 | 839 | 3,938 | 3,367 | 3,798 | 1,236 |
| cells | iPSC (AK02) | iPSC (AK02) | iPSC (AITS) | iPSC (AITS) | ESC (AITS) | EpiLC |

##### Geometrization of transcriptome data

Consider RNA-seq data generated using RNA-sequencing technology, which quantifies gene ex-pression by sequencing RNA transcripts and mapping the resulting reads to a reference genome or transcriptome. The number of reads aligned to each gene is then counted, producing a gene expression matrix in which features represent genes and entries indicate expression levels. In bulk RNA-seq data, each sample corresponds to a population of cells representing a specific cell type or condition, whereas in scRNA-seq data, each sample represents an individual cell [35, 36]. Let us denote species abstractly by s *∈{*A, B*}*, where the symbols A and B are species such as mice and humans. Suppose that 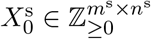 represents raw count bulk or scRNA-seq data from species s, with 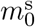 genes and 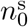 *m*_0_ *n*_0_ samples. Here, the input RNA-seq data from distinct species can have distinct gene lists and different numbers of samples.

Geometrization involves the preprocessing of input data and its conversion into a distance matrix, as introduced in the following steps.

###### Noise reduction (optional)

Although not required, we recommend applying RECODE (resolution of the curse of dimensionality) [37, 38], a cutting-edge noise reduction method, to scRNA-seq data as an initial preprocessing step. This step mitigates the curse of dimensionality, a statistical inconsistency problem caused by the accumulation of noise in high-dimensional data analysis, which leads to accurate distance computations crucial for Gromov-Wasserstein optimal transport. For bulk RNA-seq data, we skip this step. The original bulk RNA-seq data, or the denoised scRNA-seq data, is denoted by 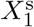.

##### Total count normalization

To adjust for sequencing depth and ensure accurate comparisons of gene expression levels, we normalize 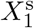 to obtain 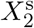, defined component-wise as

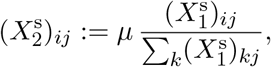

where (·)_*ij*_ denotes the (*i, j*)-entry of the matrix, and *µ* is a total normalization factor. For bulk RNA-seq data, we set *µ* = 10^6^ (reads per million) data [39]. For scRNA-seq data, we use *µ* = 10^5^ (reads per 100,000) [40].

###### Log Transformation

Assuming that the normalized data follow a log-normal distribution, we apply a logarithmic transformation to obtain 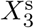, defined component-wise by 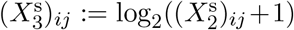. Total count normalization and log transformation steps are part of standard preprocessing procedures, and are consistent with previous studies supporting the validity of applying a logarithmic transformation to total count-normalized RNA-seq data [41, 42].

###### Filtering

To enhance the interpretability of gene-to-gene correspondences and reduce computational cost, we mask lowly expressed genes, which are commonly present in RNA-seq data, using a *gene expression cutoff parameter c*^s^ *∈*ℝ _*≥*0_, applied independently for each species. To reflect experimental cell state conditions, we define the post-cutoff matrix 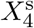 using two masking policies: *Element-wise masking* :

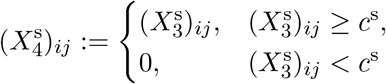

and *gene-wise masking*:

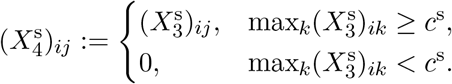

The element-wise masking sets low expression values that are indistinguishable from noise to zero, emphasizing dynamic changes in expression patterns; it is suitable for time-course data or datasets including multiple cell types. However, for static cell populations with relatively uniform transcriptome distributions, the element-wise masking may misemphasize variations of gene expression around the cutoff. Therefore, for static cell populations, gene-wise masking should be used to uniformly set lowly expressed genes to zero across all samples, avoiding selective adjustments for individual genes.

We determine the gene expression cutoff parameter *c*^s^ based on the distribution of expression values: if a local minimum exists in the distribution of maximum expression values across samples, we adopt that value; otherwise, we use a fixed percentile of the maximum expression (e.g., 5%). After masking lowly expressed genes, any gene with zero expression across all samples is removed from 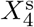, producing 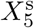.

*Gene selection* (optional): Optionally, we narrow down our scope of analysis by selecting pre-defined relevant genes, and we denote the sliced data by *X*^s^, retaining only the selected rows of genes. For example, we can select transcription factor genes, focusing on those that regulate gene expression, to study the underlying GRNs. A list of human transcription factor genes is available [43], and we can also use it to slice data of other species, based on the gene names.

Hereafter, we denote the post-preprocessed data 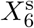as *m*^s^ *× n*^s^ matrix 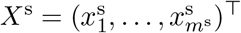. Here, 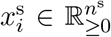 denotes the processed expression vector of *i*th gene, *m*^s^ the number of (post-selected) genes, and *n*^s^ the number of samples.

###### Distance matrix computation

Post-preprocessed RNA-seq dataset *X*^s^ defines a point cloud in *n*^s^-dimensional Euclidean space whose dimension matches its number of samples. The greater the discrepancy in dimensions of the spaces, the more pronounced the dimensionality-induced inconsistency of scales of geometries defined by usual distances becomes, especially for Euclidean distance, which tends to diverge as dimensionality increases. To reflect the similarity between point clouds with varying scales and to mitigate this distortion, we use the mean-normalized Euclidean distance

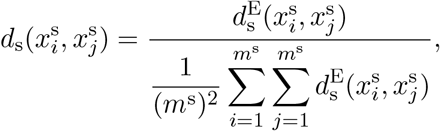

where 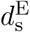 is the standard Euclidean distance given as

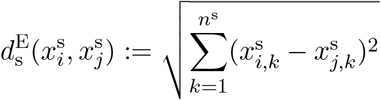

and 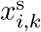 is the *k*th entry of 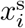. Here, the denominator reflects the mean Euclidean distance in *n*^s^-dimensional space, compensating for dimensionality effects. We compute a distance matrix 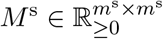, which represents the preprocessed data geometrically, by 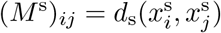.

##### Entropically regularized Gromov-Wasserstein optimal transport

We introduce the entropically regularized Gromov-Wasserstein optimal transport between geormetrizations, *M* ^A^ and *M* ^B^, for species A and B, the mathematical core of Species-OT. We define the transport polytope by

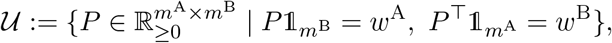

where 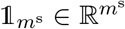 is the vector whose entries are all 1 and 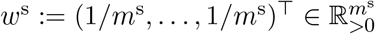, which is the homogeneous weight determined by the number of genes *m*^s^. For *p >* 0, the *p*-Gromov-Wasserstein cost associated with a coupling *P ∈𝒰* is given by

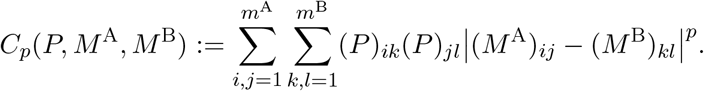

Then, we define the entropically regularized Gromov-Wasserstein optimal transport problem as the following minimization problem for a coupling *P ∈𝒰* :

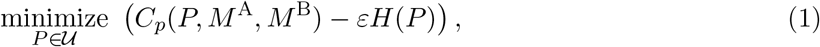

where 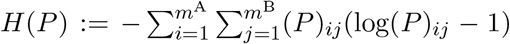 is the entropy of *P*, and *ε >* 0 is the entropic regularization parameter, which controls the strength of the convexification. A larger value of *ε* biases the solution toward a more diffused and homogeneous distribution. The solution 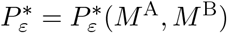 to optimization problem (1) is called the *entropic optimal transport plan*, and the associated transport cost

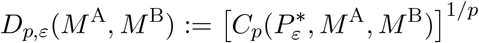

is called the *entropic Gromov-Wasserstein distance*.

If *p* = 2, the entropically regularized Gromov-Wasserstein optimal transport problem (1) reduces to a sequence of entropically regularized first-order problems, which can be solved iteratively using the Sinkhorn algorithm [44], and we can use a Gromov-Wasserstein optimal transport solver in the Python package OTT-JAX [45] to compute the numerical solution to the entropically regularized Gromov-Wasserstein optimal transport problem. Here, the numerical solution is not guaranteed to be the global minimizer of the original problem (*ε* = 0) [44], but this is the best available method at present. Hereafter, we always take *p* = 2.

In the formulation of Gromov-Wasserstein optimal transport, an inhomogeneous weight can be used in place of the homogeneous weight *w*^s^ to introduce gene-specific biases. In this study, however, we assume no such biases when computing gene-to-gene correspondences. Moreover, we can use other geometrizations, e.g., a distance matrix with Hamming distance, and further combine it with inhomogeneous weights. Accordingly, many types of omics data that are convertible to distance matrices, including proteome, genome, and epigenome data, could also be used as input data.

##### Formulation of Species-OT for a pair of transcriptome datasets

As inputs, consider a pair of raw count RNA-seq data 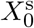 from species s *∈ {*A, B*}*, with 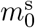 genes and 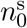 samples. Species-OT applies the geometrization to each input data and obtains distance matrices *M* ^s^ with size *m*^s^, where *m*^s^ is the number of genes after the preprocessing. Species-OT solves the entropically regularized Gromov-Wasserstein optimal transport problem (1) between distance matrices *M* ^s^, and yields the optimal transport plan 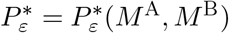 and the entropic Gromov-Wasserstein distance *D*_2,*ε*_(*M* ^A^, *M* ^B^). Regarding the entropic regularization parameter *ε*, Species-OT seeks the smallest value for which the entropically regularized Gromov-Wasserstein optimal transport problem converges numerically using the Gromov-Wasserstein optimal transport solver, as we wish to avoid too much convexification. The precise binary search-like algorithm is described in the computational parameter settings section.

Now, we transform the resulting optimal transport plan 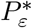 into a biologically interpretable form. The entry 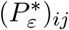 represents the probability density that the mass (weight) of the *i*th gene of species A is transported to the *j*th gene of species B. Because these values are defined so that the row and column sums match the corresponding weights, they are not directly interpretable in biological terms. To address this, we define the normalized optimal transport plan 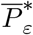 as

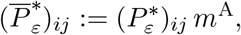

where *m*^A^ denotes the number of genes for species A. In this form, the row sums of 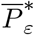 are equal to 1, and each entry 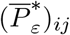 represents the percentage of mass transported from the *i*th gene of species A to the *j*th gene of species B, which can be interpreted as the probability of their correspondence.

Henceforth, we refer to the entropic Gromov-Wasserstein distance *D*_2,*ε*_(*M* ^A^, *M* ^B^) as the *transcriptomic distance*, the normalized optimal transport plan 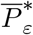 as the *gene-to-gene correspondence* from species A to B, and each entry 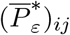 as the *corresponding rate* from the *i*th gene of species A to the *j*th gene of species B.

##### Formulation of Species-OT for multiple transcriptome datasets

As inputs of Species-OT extended to handle more than two input datasets, consider RNA-seq data from multiple species s *∈ {*S_1_, … S_*K*_*}*, where *K ∈ {*2, 3, … *}* is the number of species. Pairwise application of Species-OT produces an array of transcriptomic distances 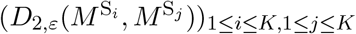, which we refer to as the *transcriptomic distance array*.

If *ε >* 0, this array is generally non-symmetric 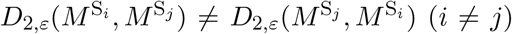 (*i* ≠ *j*), since the algorithm does not guarantee convergence to the same solution when the source and target distance matrices are swapped. In addition, the entropy regularization introduces a positive bias, resulting in a nonzero entropic Gromov-Wasserstein distance even between identical matrices, i.e., 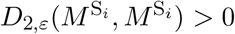. Because these characteristics hinder biological interpretability, we introduce a symmetrization procedure inspired by the Sinkhorn divergence [46]. We define the Sinkhorn-entropic Gromov-Wasserstein distance *SD*_2,*ε*_ by

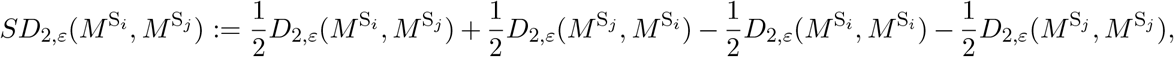

obtaining a symmetric matrix 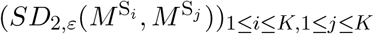, with zeros on the diagonal, whose entries are the Sinkhorn-entropic Gromov-Wasserstein distances. We refer to this matrix as the *transcriptomic discrepancy array* and its entry as *transcriptomic discrepancy*. Applying agglomerative hierarchical clustering (nearest point algorithm) to the transcriptomic discrepancy array, we obtain a dendrogram, which we refer to as *transcriptomic discrepancy tree*. The transcriptomic discrepancy array holds all pairwise quantifications of how species, or tissues from which input data are obtained, are transcriptomically similar to each other, and the transcriptomic discrepancy tree offers a concise visualization of the overall transcriptomic relationships, which can be interpreted as an extension to the phylogenetic tree at the cell-to-cell level.

#### Quantification and statistical analysis

##### Computational parameter settings

We focused on transcription factor genes among all the genes included in the input transcriptome data. We assumed that mammalian transcription factors were well-conserved, and used the same set of human transcription factor genes [43] for all the species, to select genes in the geometrization step.

The specific gene expression cutoff parameter *c*^s^ for each species in the preprocessing was determined by the distribution of the maximal expression levels in the cells. We observed the bimodality in this distribution and set the gene expression cutoff parameter at the valley between the two peaks in the distribution. For bulk RNA-seq data consisting of time-course samples, we applied element-wise masking, which evaluates expression levels in each sample independently and is suitable when samples represent distinct cell types. For scRNA-seq data, where each sample corresponds to an individual cell, we applied gene-wise masking, which retains low expression values if the gene is significantly expressed in at least one cell, and is therefore suited for data whose variable expression levels can be biologically meaningful (see Geometrization of transcriptome data section in the method details section).

The entropic regularization parameters *ε* were selected so that it convexifies the minimization problem (1) adequately, as we wish to keep the geometric structure of the original problem and make the minimization problem mathematically tractable. To ensure that the entropic regularization parameters are large enough, we first determined the smallest value with which all the pairwise entropically regularized Gromov-Wasserstein optimal transport problems converged numerically using the Gromov-Wasserstein optimal transport solver in the Python package OTT-JAX [45], with tolerance of 0.0001. We numerically sought the minimum values of the entropic regularization parameters *ε* that converge to the solution by the following binary search-like algorithm: We computed the entropically regularized Gromov-Wasserstein optimal transport problems iteratively, starting with a sufficiently large entropic regularization parameter *ε*. We took a half of the entropic regularization parameter *ε* each time we got the numerical convergence, until we got a convergence error. We then tried the intermediate value for which we got convergence and non-convergence, and we proceeded with taking half of the value if we got convergence, and tried the intermediate value for which we got convergence and non-convergence again. We repeated this iteration until the difference between the values for which we get convergence and non-convergence becomes less than the tolerance 0.0001. We then selected entropic regularization parameters larger than the minimum converging value *ε*_min_. We observed that the resulting gene-to-gene correspondences varied with the parameter in the following manner: as the parameter increased, the correspondence rates became more diffuse, while the set of top corresponding genes remained largely unchanged, with only occasional slight shifts in their ranks. Parameters used and relevant values are summarized in Tables 3–4.

**Table 3:** Parameters used and relevant values for the bulk RNA-seq data from the three mammalian species.

|  | human | macaque | mouse |
| --- | --- | --- | --- |
| gene selection criterion | transcription factor genes |  |  |
| masking policy | element-wise masking |  |  |
| # genes remained | 971 | 942 | 785 |
| cutoff parameter $c^s$ | 2.8977 | 2.8748 | 2.6878 |
| minimum converging $\varepsilon_{\min}$ | 0.001953125 | | |
| entropic parameter $\varepsilon$ | 0.01 | | |

**Table 4:** Parameters used and relevant values for the scRNA-seq data from the six mammalian species.

|  | human | chimpanzee | gorilla | orangutan | macaque | mouse |
| --- | --- | --- | --- | --- | --- | --- |
| gene selection criterion | transcription factor genes |  |  |  |  |  |
| masking policy | gene-wise masking |  |  |  |  |  |
| # genes remained | 1,144 | 1,083 | 1,026 | 1,107 | 1,129 | 634 |
| cutoff parameter $c^s$ | 1.2861 | 1.3055 | 1.2201 | 1.2023 | 1.0464 | 1.0827 |
| minimum converging $\varepsilon_{\min}$ | 0.00693359375 | | | | | |
| entropic parameter $\varepsilon$ | 0.1 | | | | | |

##### Differential expression analysis and selection of Wnt-related genes

To identify genes whose expression was significantly downregulated in cells prepared using culture condition AITS compared to AK02 (Figure 5D), we performed a differential expression analysis between the two conditions. For each gene, normalized log-transformed expression values in AK02 and AITS cells were compared using Welch’s *t* -test, followed by multiple testing correction. We then focused on genes involved in the Wnt signaling pathway, selecting seven genes: *CXXC4, PPARA, SIX3, SOX9, SREBF1, RARG*, and *RORA*. To assess how their expression varied among species, we extracted normalized expression values for each species in the dataset and plotted them as boxplots. Expression values were *z* -score transformed on a per-gene basis to enable direct comparison of relative changes across species while accounting for gene-specific dynamic ranges.

#### Experimental model and subject details

##### Ethics

All experimental procedures involving human iPSCs were approved by the Institutional Review Board of Kyoto University and conducted in accordance with the guidelines of the Ministry of Education, Culture, Sports, Science and Technology of Japan. Experimental procedures involving great ape iPSCs were approved by the Animal Care and Use Committee of the Kyoto University Primate Research Institute (KUPRI) and the Center for the Evolutionary Origins of Human Behavior (EHUB), and were performed in accordance with the Guidelines for the Care and Use of Nonhuman Primates (3rd edition, 2010), published by KUPRI. This study did not involve any experiments using live animals.

##### Cell lines

Human iPSCs (1390G3, 1390G3–AGVT [bearing *TFAP2C* –*EGFP* and *VASA*–*tdTomato* transgenes]), mouse ESC (BDF1-2-1-BVSC [bearing the *Blimp1* –*mVenus* and *Stella*–*ECFP* transgenes]), cynomolgus monkey ESCs (15XRi, 15XRi-AGVT) and chimpanzee iPSCs (0274F-2, Mari4) were cultured as previously described [11, 34, 47, 48, 49, 50, 51, 52, 53].

Briefly, human and chimpanzee iPSCs were cultured in StemFit AK02N (Ajinomoto) on plastic plates coated with iMatrix (iMATRIX-511; Nippi, 892014).

Mouse ESCs were maintained in N2B27 medium supplemented with 0.4 µM PD0325901 (Stemgent, 04-2006), 3 µM CHIR99021 (Bio Vision, 4423), and 1000 U/mL leukemia inhibitory factor (LIF; Millipore, ESG1107) on plastic plates coated with 0.01% poly-L-ornithine (Sigma Aldrich, P3655) and 100 ng/mL laminin (BD Biosciences, 354232).

Cynomolgus monkey ESCs, as well as gorilla iPSCs (Wif131; established as described below) and orangutan iPSCs (TobaK; gifted by Dr. Keisuke Okita), were cultured in AITS-IF20 medium [47, 48]. AITS-IF20 consists of a 1:1 mixture of Advanced RPMI 1640 (Thermo Fisher Scientific, 12633012), and Neurobasal medium (Thermo Fisher Scientific, 12348017), supplemented with 1.6% (w/v) Albumax (Thermo Fisher Scientific, 11020062), 1 × ITS (Thermo Fisher Scientific, 41400045), 1% non-essential amino acids (NEAA; Thermo Fisher Scientific, 11140-050), 2 mM L-glutamine (Thermo Fisher Scientific, 25030081), 25 U/mL penicillin/streptomycin (Thermo Fisher Scientific, 15140148), 20 ng/mL bFGF (Wako, 060-04543), and 2.5 µM IWR-1 (Sigma Aldrich, IO161) and was used on mouse embryonic fibroblasts (MEFs).

##### Establishment of gorilla iPSCs

Gorilla dermal fibroblasts from skin specimens (Western Lowland Gorilla, male, 45-year-old, GAIN-ID: 0023, Willie) were obtained from the Great Ape Information Network (GAIN, Japan) and reprogrammed by introducing the Yamanaka four factors using the Sendai virus transduction method (CytoTune-iPS 2.0; Nacalai Tesque), according to the manufacturer’s instructions. On day 5 after transduction, the cells were replated on MEFs, and the resultant colonies were picked on day 18. After establishment, the iPSCs were cultured in AITS-IF20 medium as described above.

##### Human and cynomolgus macaque PGCLC induction

Human PGCLC induction via iMeLC was performed essentially as previously described [11, 51]. To induce iMeLCs, iPSCs were cultured for approximately 48 hours in iMeLC induction medium consisting of GMEM supplemented with 15% KSR, 1% NEAA, 0.1 mg/mL streptomycin, 2 mM L-glutamine, 2 mM sodium pyruvate, and 0.1 mM *β*-mercaptoethanol. The medium was further supplemented with 3 µM CHIR99021, 50 ng/mL Activin A, and 10 µM Y27632 (Tocris, 1254), and cells were cultured on fibronectin-coated plates (Millipore, FC010). To induce PGCLCs, dissociated iMeLCs were plated in PGCLC induction medium consisting of either GMEM or Advanced RPMI 1640, supplemented with 15% KSR, 1% NEAA, 1% penicillin-streptomycin, 2 mM L-glutamine, 2 mM sodium pyruvate, and 0.1 mM *β*-mercaptoethanol. The medium also contained 200 ng/mL BMP4, 1000 U/mL LIF, 100 ng/mL SCF, 50 ng/mL EGF, and 10 µM Y27632. Cultures were maintained in V-bottom 96-well plates (Greiner, 651970).

Cynomolgus monkey PGCLC induction was performed as previously described [47, 48]. CyESCs were plated into wells of low-binding V-bottom 96-well plates in aRB27 medium, which consists of Advanced RPMI 1640 supplemented with 2× B27 without vitamin A (Gibco, 12587-001), 0.1 mM NEAA, 2 mM L-glutamine, and 25 U/mL penicillin-streptomycin. The medium was further supplemented with 200 ng/mL BMP4, 1000 U/mL LIF, 100 ng/mL SCF, 50 ng/mL human EGF, and 10 µM Y27632.

##### Cell collection

For the collection of ESCs, iMeLCs, and EpiLCs, cells were dissociated and suspended in 100 µL of FACS buffer (0.1% BSA Fraction V (Thermo Fisher Scientific, 15260037) in PBS) supplemented with 10 µM Y27632. For feeder-dependent cultures, cells were incubated on ice for 30 minutes with 5 µL APC-conjugated anti-mouse/rat CD29 antibodies (Biolegend, 102216) to exclude MEF feeder cells. After washing once with 1 mL FACS buffer, cells were resuspended in 500 µL FACS buffer containing 10 µM Y27632 and 5 µg/mL propidium iodide (PI; Sigma Aldrich, P4170). The cell suspension was filtered through a cell strainer, and CD29/PI double-negative cells were collected using a flow cytometer (FACS Aria III; BD Biosciences) according to the manufacturer’s instructions.

For human and cynomolgus monkey PGCLCs, floating aggregates were treated with 300 µL of 1 mg/mL type IV collagenase (MPBio, 195110) in aRB27 supplemented with 10 µM Y27632 at 37°C for 15 minutes. After washing twice with 1 mL PBS, cells were dissociated in 300 µL of 0.25% Trypsin/EDTA containing 10 µM Y27632 at 37°C for 15 minutes. The cells were filtered through a cell strainer, and AG-reporter-positive cells were collected using a flow cytometer (FACS Aria III; BD Biosciences) according to the manufacturer’s instructions.

##### Bulk 3’RNA-seq data acquisition

cDNA library construction from 1 ng of total RNA was performed as previously described [48, 54, 55]. The quality and concentration of the resulting library DNA were evaluated using a LabChip GX system (Perkin Elmer), a Qubit dsDNA HS Assay Kit (Thermo Fisher Scientific, Q32854) and TaqMan PCR using THUNDERBIRD Probe qPCR Mix (TOYOBO, QPS-101) and a TaqMan probe (Applied Biosystems, Ac04364396). Sequencing was performed using a NextSeq 500 High Output Kit v2.5 (75 cycles; Illumina).

##### 10X scRNA-seq data acquisition

ScRNA-seq libraries of 10X data were generated using the 10X Genomics Chromium Controller (10X Genomics) and Chromium Single Cell 3’ Reagent Kits v3.1 according to the manufacturer’s instructions. For mouse EpiLC, the cell suspension was subjected to Cell Multiplexing Oligo labeling according to the manufacturer’s instructions. For reverse transcription, cDNA amplification, and sample indexing were performed using an Eppendorf Mastercycler. The final libraries were quantified using a KAPA library quantification kit (KAPA Biosystems, KK4824), and the fragment size distribution of the libraries was determined using a LabChip GX DNA high sensitivity kit (Perkin Elmer). Pooled libraries were then sequenced using a NovaSeq 6000 platform with an SP or S1 100-cycle kit (Illumina).

##### Mapping of transcriptome reads and conversion to gene expression levels

Genome sequences (`GRCm38` for mouse, `GRCh38p12` for human, `Clint_PTRv2/panTro6` for chimpanzee, `gorGor6` for gorilla, `Susie_PABv2/ponAbe3` for orangutan, and `macFasRKS1912v2` for cynomolgus monkey) were obtained from the NCBI FTP site. Gene annotations for human were downloaded from GENCODE v38 “Comprehensive”. Genes located on reference chromosomes and annotated with `gene_types` other than `pseudogene, tRNA, rRNA, scRNA, snRNA, snoRNA, vault_RNA, ribozyme,` and `TEC` were retained for downstream analysis. These filtered annotations are referred to simply as GENCODE hereafter. Additionally, genes that could be converted from GENCODE using LiftOver and validated with LiftOff were incorporated to improve gene annotations for non-human primates.

For bulk 3’ RNA-seq data, reads were processed using Cutadapt v1.18 and mapped using TopHat2 v2.1.1 with Bowtie2 v2.3.4.1, as described previously [48, 54, 55]. Since 3’ RNA-seq captures only the very 3’ end of transcripts, expression levels were quantified at the gene level (Entrez gene) rather than the mRNA isoform level.

For 10X scRNA-seq data, reads were processed using Cell Ranger v6.0.1 without including intronic reads.

##### Multiplet Identification

In 10X scRNA-seq data, low-quality cells were defined as those with fewer than 10,000 UMIs, fewer than 2,000 detected genes, or more than 20% of reads mapped to mitochondrial genes. These low-quality cells, along with putative multiplets, were identified using the Scrublet Python package (v0.2.3) [56]. The resulting filtered data was then uploaded as supplementary data and used as the scRNA-seq data (Table 2) for input to Species-OT.

##### Processing of bulk RNA-seq data

We processed mouse bulk RNA-seq data (GSE67259) to generate an expression matrix indexed by human gene symbols. A human gene dictionary was assembled from Ensembl and HGNC annotations, integrated with human–mouse orthology from BioMart, and linked to mouse Entrez IDs via MGI coordinates. These resources were merged into a cross-species mapping table, with many-to-many relationships simplified and duplicates removed. Using this dictionary, we re-annotated the mouse dataset by excluding non-gene entries, relabeling rows with human symbols, and collapsing duplicates. The resulting table preserves the structure of the original data while enhancing cross-species analysis. We used human and macaque bulk RNA-seq data (GSE307647) to generate expression matrices indexed by human gene symbols. The resulting processed data was then uploaded as supplementary data and used as the bulk RNA-seq data (Table 1) for input to Species-OT.

## References

[1] International Human Genome Sequencing Consortium. “Initial sequencing and analysis of the human genome”. Nature 409 (2001), pp. 860–921. doi: 10.1038/35057062.

[2] The Chimpanzee Sequencing and Analysis Consortium. “Initial sequence of the chimpanzee genome and comparison with the human genome”. Nature 437 (2005), pp. 69–87. doi: 10.1038/nature04072.

[3] Mouse Genome Sequencing Consortium. “Initial sequencing and comparative analysis of the mouse genome”. Nature 420 (2002), pp. 520–562. doi: 10.1038/nature01262.

[4] W. M. Fitch. “Distinguishing Homologous from Analogous Proteins”. Systematic Biology 19.2 (1970), pp. 99–113. doi: 10.2307/2412448.

[5] E. L. L. Sonnhammer and E. V. Koonin. “Orthology, paralogy and proposed classification for paralog subtypes”. Trends in Genetics 18.12 (2002), pp. 619–620. doi: 10.1016/S0168-9525(02)02793-2.

[6] T. Gabaldón and E. V. Koonin. “Functional and evolutionary implications of gene orthology”. Nature Reviews Genetics 14.5 (2013), pp. 360–366. doi: 10.1038/nrg3456.

[7] D. Villar, C. Berthelot, S. Aldridge, et al. “Enhancer Evolution across 20 Mammalian Species”. Cell 160.3 (2015), pp. 554–566. doi: 10.1016/j.cell.2015.01.006.

[8] G. Bourque, K. H. Burns, M. Gehring, et al. “Ten things you should know about transposable elements”. Genome Biology 19.1 (2018), p. 199. doi: 10.1186/s13059-018-1577-z.

[9] J. Nichols and A. Smith. “Naive and Primed Pluripotent States”. Cell Stem Cell 4.6 (2009), pp. 487–492. doi: 10.1016/j.stem.2009.05.015.

[10] K. Semi and Y. Takashima. “Pluripotent stem cells for the study of early human embryology”. Development, Growth & Differentiation 63.2 (2021), pp. 104–115. doi: 10.1111/dgd.12715.

[11] Y. Kojima, K. Sasaki, S. Yokobayashi, et al. “Evolutionarily Distinctive Transcriptional and Signaling Programs Drive Human Germ Cell Lineage Specification from Pluripotent Stem Cells”. Cell Stem Cell 21.4 (2017), pp. 517–532. doi: 10.1016/j.stem.2017.09.005.

[12] K. Biharie, L. Michielsen, M. J. T. Reinders, et al. “Cell type matching across species using protein embeddings and transfer learning”. Bioinformatics 39.Supplement_1 (2023), pp. i404–i412. doi: 10.1093/bioinformatics/btad248.

[13] A. J. Tarashansky, J. M. Musser, M. Khariton, et al. “Mapping single-cell atlases throughout Metazoa unravels cell type evolution”. eLife 10 (2021), e66747. doi: 10.7554/eLife.66747.

[14] M. M. Saint-Antoine and A. Singh. “Network inference in systems biology: recent developments, challenges, and applications”. Current Opinion in Biotechnology 63 (2020), pp. 89–98. doi: 10.1016/j.copbio.2019.12.002.

[15] G. Schiebinger, J. Shu, M. Tabaka, et al. “Optimal-Transport Analysis of Single-Cell Gene Expression Identifies Developmental Trajectories in Reprogramming”. Cell 176.4 (2019), pp. 928–943. doi: 10.1016/j.cell.2019.01.006.

[16] M. Nitzan, N. Karaiskos, N. Friedman, et al. “Gene expression cartography”. Nature 576 (2019), pp. 132–137. doi: 10.1038/s41586-019-1773-3.

[17] K. Cao, Y. Hong, and L. Wan. “Manifold alignment for heterogeneous single-cell multi-omics data integration using Pamona”. Bioinformatics 38.1 (2021), pp. 211–219. doi: 10.1093/bioinformatics/btab594.

[18] P. Demetci, R. Santorella, B. Sandstede, et al. “SCOT: Single-Cell Multi-Omics Alignment with Optimal Transport”. Journal of Computational Biology 29.1 (2022), pp. 3–18. doi: 10.1089/cmb.2021.0446.

[19] M. Saitou and K. Hayashi. “Mammalian in vitro gametogenesis”. Science 374.6563 (2021), pp. 1–9. doi: 10.1126/science.aaz6830.

[20] Y. Kojima, C. Yamashiro, Y. Murase, et al. “GATA transcription factors, SOX17 and TFAP2C, drive the human germ-cell specification program”. Life Science Alliance 4.5 (2021). doi: 10.26508/lsa.202000974.

[21] A. D. Yates, P. Achuthan, W. Akanni, et al. “Ensembl 2020”. Nucleic Acids Research 48.D1 (2020), pp. D682–D688. doi: 10.1093/nar/gkz966.

[22] N. Irie, L. Weinberger, W. W. Tang, et al. “SOX17 Is a Critical Specifier of Human Primordial Germ Cell Fate”. Cell 160.1 (2015), pp. 253–268. doi: 10.1016/j.cell.2014.12.013.

[23] Y. L. Yamaguchi, S. S. Tanaka, M. Kumagai, et al. “Sall4 Is Essential for Mouse Primordial Germ Cell Specification by Suppressing Somatic Cell Program Genes”. Stem Cells 33.1 (2015), pp. 289–300. doi: 10.1002/stem.1853.

[24] T. Nakamura, K. Fujiwara, M. Saitou, et al. “Non-human primates as a model for human development”. Stem Cell Reports 16.5 (2021), pp. 1093–1103. doi: 10.1016/j.stemcr.2021.03.021.

[25] P. Perelman, W. E. Johnson, C. Roos, et al. “A Molecular Phylogeny of Living Primates”. PLOS Genetics 7.3 (2011), e1001342. doi: 10.1371/journal.pgen.1001342.

[26] Y. Zheng, X. Xue, Y. Shao, et al. “Controlled modelling of human epiblast and amnion development using stem cells”. Nature 573.7774 (2019), pp. 421–425. doi: 10.1038/s41586-019-1535-2.

[27] K. Jo, S. Teague, B. Chen, et al. “Efficient differentiation of human primordial germ cells through geometric control reveals a key role for Nodal signaling”. eLife 11 (2022), e72811. doi: 10.7554/eLife.72811.

[28] L. Yu, Y. Wei, H.-X. Sun, et al. “Derivation of Intermediate Pluripotent Stem Cells Amenable to Primordial Germ Cell Specification”. Cell Stem Cell 28.3 (2021), 550–567.e12. doi: 10.1016/j.stem.2020.11.003.

[29] J. D. Buenrostro, P. G. Giresi, L. C. Zaba, et al. “Transposition of native chromatin for fast and sensitive epigenomic profiling of open chromatin, DNA-binding proteins and nucleosome position”. Nature Methods 10.12 (2013), pp. 1213–1218. doi: 10.1038/nmeth.2688.

[30] D. S. Johnson, A. Mortazavi, R. M. Myers, et al. “Genome-Wide Mapping of in Vivo Protein-DNA Interactions”. Science 316.5830 (2007), pp. 1497–1502. doi: 10.1126/science.1141319.

[31] E. Lieberman-Aiden, N. L. van Berkum, L. Williams, et al. “Comprehensive Mapping of Long-Range Interactions Reveals Folding Principles of the Human Genome”. Science 326.5950 (2009), pp. 289–293. doi: 10.1126/science.1181369.

[32] R. Aebersold and M. Mann. “Mass spectrometry-based proteomics”. Nature 422.6928 (2003), pp. 198–207. doi: 10.1038/nature01511.

[33] K. Hayashi and M. Saitou. “Generation of eggs from mouse embryonic stem cells and induced pluripotent stem cells”. Nature Protocols 8.8 (2013), pp. 1513–1524. doi: 10.1038/nprot.2013.090.

[34] M. Nakagawa, Y. Taniguchi, S. Senda, et al. “A novel efficient feeder-free culture system for the derivation of human induced pluripotent stem cells”. Scientific Reports 4.1 (2014), p. 3594. doi: 10.1038/srep03594.

[35] Z. Wang, M. Gerstein, and M. Snyder. “RNA-Seq: a revolutionary tool for transcriptomics”. Nature Reviews Genetics 10 (2009), pp. 57–63. doi: 10.1038/nrg2484.

[36] F. Tang, C. Barbacioru, Y. Wang, et al. “mRNA-Seq whole-transcriptome analysis of a single cell”. Nature Methods 6 (2009), pp. 377–382. doi: 10.1038/nmeth.1315.

[37] Y. Imoto, T. Nakamura, E. G. Escolar, et al. “Resolution of the curse of dimensionality in single-cell RNA sequencing data analysis”. Life Science Alliance 5.12 (2022), pp. 1–23. doi: 10.26508/lsa.202201591.

[38] Y. Imoto. “Comprehensive noise reduction in single-cell data with the RECODE platform”. Cell Reports Methods 5.10 (2025), p. 101178. doi: 10.1016/j.crmeth.2025.101178.

[39] T. Nakamura, I. Okamoto, K. Sasaki, et al. “A developmental coordinate of pluripotency among mice, monkeys and humans”. Nature 537 (2016), pp. 57–62. doi: 10.1038/nature19096.

[40] Y. Imoto. “Accurate highly variable gene selection using RECODE in scRNA-seq data analysis”. bioRxiv (2025). Preprint, p. 2025.06.23.661026. doi: 10.1101/2025.06.23.661026.

[41] I. Zwiener, B. Frisch, and H. Binder. “Transforming RNA-Seq Data to Improve the Performance of Prognostic Gene Signatures”. PLOS ONE 9.1 (2014), e85150. doi: 10.1371/journal.pone.0085150.

[42] M. Gierliński, C. Cole, P. Schofield, et al. “Statistical models for RNA-seq data derived from a two-condition 48-replicate experiment”. Bioinformatics 31.22 (2015), pp. 3625–3630. doi: 10.1093/bioinformatics/btv425.

[43] S. A. Lambert, A. Jolma, L. F. Campitelli, et al. “The Human Transcription Factors”. Cell 172.4 (2018), pp. 650–665. doi: 10.1016/j.cell.2018.09.045.

[44] J. Solomon, G. Peyré, V. G. Kim, et al. “Entropic metric alignment for correspondence problems”. ACM Trans. Graph. 35.4 (2016), pp. 1–13. doi: 10.1145/2897824.2925903.

[45] M. Cuturi, L. Meng-Papaxanthos, Y. Tian, et al. Optimal Transport Tools (OTT): A JAX Toolbox for all things Wasserstein. 2022. doi: 10.48550/arXiv.2201.12324.

[46] J. Feydy, T. Séjourné, F.-X. Vialard, et al. “Interpolating between Optimal Transport and MMD using Sinkhorn Divergences”. Proc. 22nd Int. Conf. Artif. Intell. Stat. (AISTATS). Vol. 89. Proc. Mach. Learn. Res. PMLR, 2019, pp. 2681–2690. url: https://proceedings.mlr.press/v89/feydy19a.html.

[47] Y. Sakai, T. Nakamura, I. Okamoto, et al. “Induction of the germ cell fate from pluripotent stem cells in cynomolgus monkeys”. Biology of Reproduction 102.3 (2019), pp. 620–638. doi: 10.1093/biolre/ioz205.

[48] S. Gyobu-Motani, Y. Yabuta, K. Mizuta, et al. “Induction of fetal meiotic oocytes from embryonic stem cells in cynomolgus monkeys”. The EMBO Journal 42.9 (2023), pp. 1–30. doi: 10.15252/embj.2022112962.

[49] M. Nagano, B. Hu, S. Yokobayashi, et al. “Nucleome programming is required for the foundation of totipotency in mammalian germline development”. The EMBO Journal 41.13 (2022), pp. 1–34. doi: 10.15252/embj.2022110600.

[50] Y. Ishikura, H. Ohta, T. Sato, et al. “In vitro reconstitution of the whole male germ-cell development from mouse pluripotent stem cells”. Cell Stem Cell 28.12 (2021), pp. 2167–2179. doi: 10.1016/j.stem.2021.08.005.

[51] K. Sasaki, S. Yokobayashi, T. Nakamura, et al. “Robust In Vitro Induction of Human Germ Cell Fate from Pluripotent Stem Cells”. Cell Stem Cell 17.2 (2015), pp. 178–194. doi: 10.1016/j.stem.2015.06.014.

[52] R. Kitajima, R. Nakai, T. Imamura, et al. “Modeling of early neural development in vitro by direct neurosphere formation culture of chimpanzee induced pluripotent stem cells”. Stem Cell Research 44 (2020), pp. 1–13. doi: 10.1016/j.scr.2020.101749.

[53] C. Yamashiro, K. Sasaki, Y. Yabuta, et al. “Generation of human oogonia from induced pluripotent stem cells in vitro”. Science 362.6412 (2018), pp. 356–360. doi: 10.1126/science.aat1674.

[54] T. Nakamura, Y. Yabuta, I. Okamoto, et al. “SC3-seq: a method for highly parallel and quantitative measurement of single-cell gene expression”. Nucleic Acids Research 43.9 (2015), pp. 1–17. doi: 10.1093/nar/gkv134.

[55] Y. Ishikura, Y. Yabuta, H. Ohta, et al. “In Vitro Derivation and Propagation of Spermatogonial Stem Cell Activity from Mouse Pluripotent Stem Cells”. Cell Reports 17.10 (2016), pp. 2789–2804. doi: 10.1016/j.celrep.2016.11.026.

[56] S. L. Wolock, R. Lopez, and A. M. Klein. “Scrublet: Computational Identification of Cell Doublets in Single-Cell Transcriptomic Data”. Cell Systems 8.4 (2019), pp. 281–291. doi: 10.1016/j.cels.2018.11.005.

